# Viral expression of a SERCA2a-activating PLB mutant improves calcium cycling and synchronicity in dilated cardiomyopathic hiPSC-CMs

**DOI:** 10.1101/699975

**Authors:** Daniel R. Stroik, Delaine K. Ceholski, Justyna Mleczko, Paul F. Thanel, Philip A. Bidwell, Joseph M. Autry, Razvan L. Cornea, David D. Thomas

## Abstract

There is increasing momentum toward the development of gene therapy for heart failure (HF), cardiomyopathy, and other progressive cardiac diseases that correlate with impaired calcium (Ca^2+^) transport and reduced contractility. We have used FRET between fluorescently-tagged SERCA2a (the cardiac Ca^2+^ pump) and PLB (its ventricular peptide inhibitor) to test directly the effectiveness of loss-of-inhibition/gain-of-binding (LOI/GOB) PLB mutants (PLB_M_) that were engineered to compete with the binding of inhibitory wild type PLB (PLB_WT_). Our therapeutic strategy is to relieve PLB_WT_ inhibition of SERCA2a by utilizing the reserve adrenergic capacity of PLB to enhance baseline cardiac contractility. Using a FRET assay, we determined that the combination of a LOI PLB mutation (L31A) and a GOB PLB mutation (I40A) results in a novel engineered LOI/GOB PLB_M_ (L31A/I40A) that effectively competes with PLB_WT_ binding to cardiac SERCA2a in HEK293-6E cells. We demonstrated that co-expression of L31A/I40A-PLB_M_ enhances SERCA Ca-ATPase activity by increasing enzyme Ca^2+^ affinity (1/K_Ca_) in PLB_WT_-inhibited HEK cell homogenates. For an initial assessment of PLB_M_ physiological effectiveness, we used human induced pluripotent stem cell derived cardiomyocytes (hiPSC-CMs) from a healthy individual. In this system, we observed that adeno-associated virus 2 (rAAV2)-driven expression of L31A/I40A-PLB_M_ enhances the amplitude of SR Ca^2+^ release and the rate of SR Ca^2+^ re-uptake. To assess therapeutic potential, we used an hiPSC-CM model of dilated cardiomyopathy (DCM) containing PLB mutation R14del, where we observed that rAAV2-driven expression of L31A/I40A-PLB_M_ rescues arrhythmic Ca^2+^ transients and alleviates decreased Ca^2+^ transport. Based on these results, PLB_M_ transgene expression is a promising gene therapy strategy for cardiomyopathies associated with impaired Ca^2+^ transport and decreased contractility.

## 1. Introduction

Heart failure (HF) is a pathology characterized by the impaired capacity of ventricles to fill with or eject blood, constituting a leading cause of morbidity and mortality worldwide.^1^ Over the past decades, researchers have made significant progress in identifying intracellular and molecular mechanisms that are altered during HF progression. Defective sarcoplasmic reticulum (SR) Ca^2+^ transport has been identified as a potential determinant of decreased contractility in the failing hearts.^2–4^ Ca^2+^ transport from the myosol to the SR lumen in human cardiomyocytes (CM) is largely performed (~70% of Ca^2+^ ions transported) via the SR Ca-ATPase (SERCA2a).^5^ Despite recent advances in device and pharmacological therapies, rates of heart disease continue to rise due to an aging and increasingly obese population, so new treatment options are urgently needed. Increasing knowledge of the molecular mechanisms fundamental to cardiac function and disease – including HF – has expanded the therapeutic potential to target key players involved in Ca^2+^ handling, including SERCA2a.

The activity of SERCA2a is regulated by phospholamban (PLB), a 52-residue single-pass transmembrane protein expressed in the SR of cardiac muscle. PLB binds to SERCA2a and reduces its Ca^2+^ affinity, as measured from SERCA activity.^6^ PLB is in equilibrium between homopentamers and monomers, where the oligomeric state is proposed to function as a reservoir.^7^ Previous studies have identified two functional regions on the transmembrane (TM) helix of PLB.^8–10^ Single-residue mutations on one side of the TM helix diminish inhibitory function without significantly affecting PLB oligomerization state(s), whereas mutations on the other side of the TM helix modulate PLB oligomerization and can lead to enhanced SERCA inhibition.^8,11^ The regulation of gene expression has been investigated in healthy individuals and HF patients, and decreased SERCA-to-PLB ratio is associated with deteriorated cardiac function, indicating that SERCA/PLB stoichiometry and SERCA2a activity are potential therapeutic targets.^12–14^ To this end, SERCA2a transgene expression via recombinant adeno-associated virus (rAAV) was achieved in HF animal models to increase the SERCA-to-PLB ratio in the heart. Exogenous SERCA2a expression significantly enhanced cardiac function in disease models by multiple metrics.^15,16^ Recently, gene therapy designed to increase SERCA2a expression in the human heart underwent clinical trials in patients diagnosed with end-stage HF.^17^ Despite positive preliminary results in Phase 1/2a,^18^ the trial did not meet its primary end goals in phase 2b, and this has been ascribed to dosage constraints.^19^ One likely factor is the large size of SERCA2a, which limits steady-state expression levels of the gene therapy construct. Other promising strategies to increase the SERCA-to-PLB ratio include expression of miRNAs that directly target PLB expression^20–22^ and oligonucleotide-based drugs^23,24^, although these strategies have not yet produced an effective therapeutic agent.

In the present study, we have pursued an alternative approach to activate SERCA2a, by expressing a loss-of-inhibition (LOI) PLB mutant (PLB_M_) that is engineered to compete with the inhibitory wild type PLB (PLB_WT_) for SERCA2a binding (Fig. 1A) via viral delivery. In this approach, the ratio between endogenous SERCA and PLB is not targeted, rather the exogenous PLB_M_ relieves SERCA2a inhibition and enhances its Ca^2+^ transport function. Thus, this strategy is designed to overcome limitations associated with SERCA2a overexpression-based gene therapy.

**Fig. 1.**
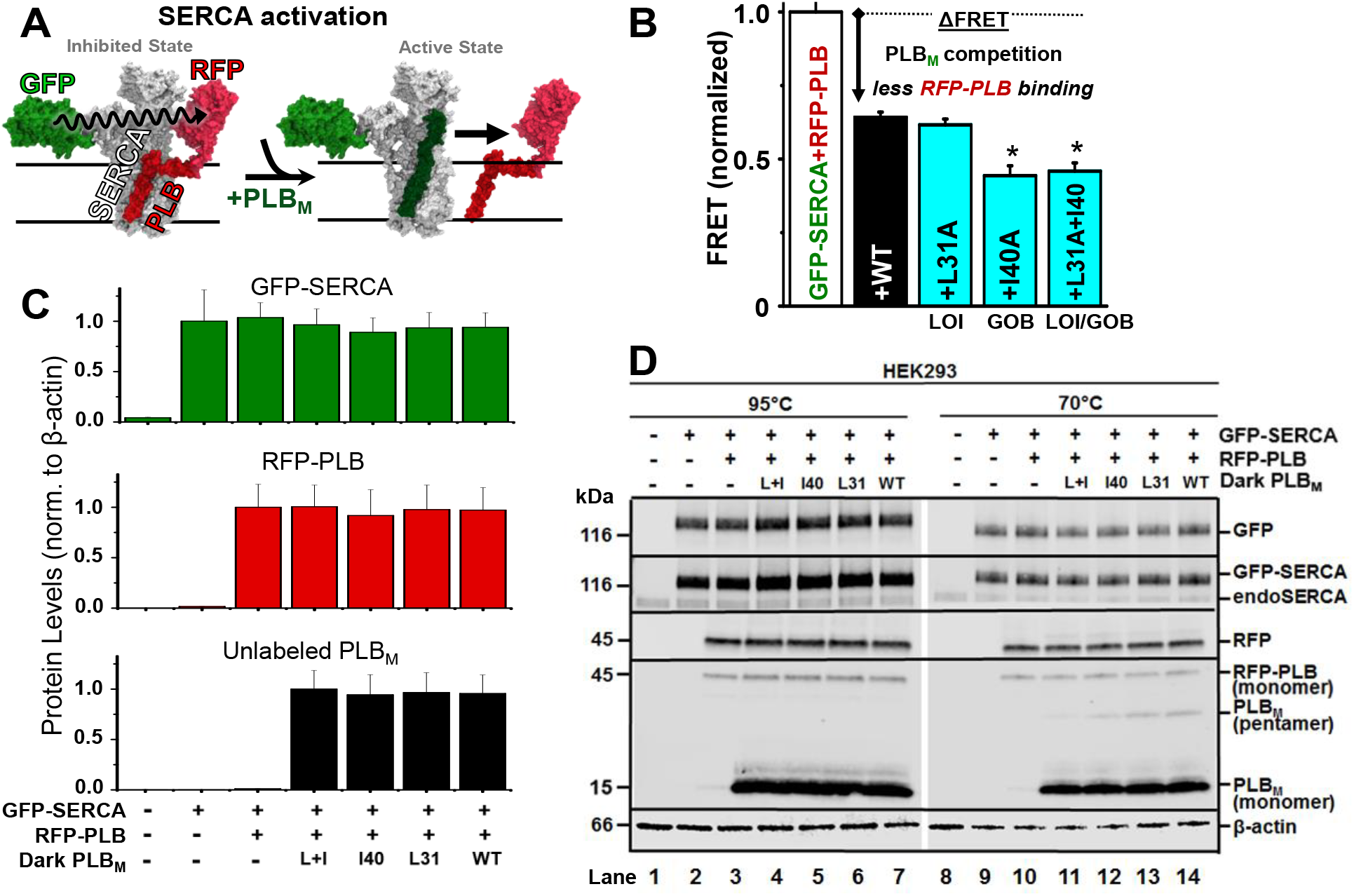
Co-expression of SERCA, PLB, and PLB_M_. (A) Schematic of gene therapy strategy: a tight-binding, non-inhibitory PLB mutant (PLB_M_, dark green) displaces inhibitory RFP-PLB (PLB_WT_, dark red), resulting in a FRET decrease. (B) FRET assay of PLB_M_ competition in HEK293-6E. GFP-SERCA-RFP-PLB_WT_ FRET (binding, *white bar*) is decreased (arrow) by PLB variants. LOI = Loss-of-inhibitory function mutant. GOB = Gain-of-inhibition mutant. (C/D) Immunoblots of HEK homogenate samples incubated at 95°C (disrupting PLB pentamer, lanes 1-7) or at 70°C (lanes 8‒14) from untransfected HEK293-6E cells (lane 1&8), cells expressing GFP-SERCA2a (lanes 2&9), or cells expressing GFP-SERCA2a+RFP-PLB (lanes 3‒7&11‒14) +L31A/I40A-PLB (lanes 4&11), +I40A-PLB (lanes 5&12), +L31A-PLB (lanes 6&13), and +WT-PLB (lanes 7&14). Primary antibodies used (from top to bottom of 1D panels) are anti-GFP, anti-SERCA2, anti-RFP, anti-PLB, and anti-β-actin. **Expression levels were quantitated by** densitometry in lanes 1‒7. Error bars indicate SEM (n=3). *P < *0.05.* **Note to journal: this is a 2-column wide figure**.

## 2. Materials and methods

### 2.1. Molecular biology

eGFP and tagRFP were fused to the N-terminus of human SERCA2a and human PLB, respectively. We have demonstrated that attachment of the fluorescent proteins at these sites does not interfere with SERCA activity or PLB inhibition.^25,26^ PLB cDNA mutations were introduced using the QuikChange mutagenesis kit (Agilent Technologies, Santa Clara, CA), and all expression plasmids were sequenced for confirmation.

### 2.2. Cell culture

Human embryonic kidney cells 293 (HEK293-6E, NRC, Canada) were cultured in FreeStyle F17 expression medium (Thermo Fisher Scientific, Waltham, MA) supplemented with 2 mM L-glutamine (Invitrogen, Waltham, MA). Cell cultures were maintained in a circular shaker (125 rpm, 37°C, 5% CO_2_ (Forma Series II Water Jacket CO_2_ Incubator, Thermo Fisher Scientific, Waltham, MA). For displacement assays, HEK293-6E cells were transiently transfected using 293fectamine with GFP-SERCA2a, RFP-PLBWT, and PLB_M_ or empty vector in a 1:7:7 molar ratio. Cells were then assayed 48 hours post-transfection.

The control hiPSC line (SKiPS-31.3) was cultured as previously described.^27^ The R14del hiPSC line was derived from the SKiPS-31.3 line using homologous recombination via CRISPR/Cas9. Monolayer cardiac differentiation was performed as described,^27^ yielding beating cardiomyocytes within 7-10 days.

### 2.3. Gene expression

RNA was extracted from uninfected and rAAV-infected hiPSC-CMs at day 37, and cDNA was synthesized as previously described.^27^ Gene expression compared to the housekeeping gene β2-microglobulin (B2M) was determined using qRT-PCR, as assessed by ΔΔC_t_ analysis. See Supplemental Table 1 for the list of primers used for qRT-PCR.

### 2.4. Immunoblot analysis

Samples were separated on a 4-20% polyacrylamide gradient gel (Bio-Rad, Hercules, CA) and transferred to polyvinylidene difluoride (PVDF) membrane. The PVDF membrane was blocked in Odyssey Blocking Buffer (LI-COR Biosciences, Lincoln, NE) followed by overnight incubation at 4 °C with the primary target antibody: rabbit anti-GFP polyclonal antibody (pAb) (1:1000; ab290, Abcam, Cambridge, United Kingdom), mouse anti-SERCA2 monoclonal antibody (mAb) (1:1000; 2A7-A1, Abcam), rabbit anti-tagRFP pAb (1:1000; ab233, Evrogen), mouse anti-PLB mAb (1:1000, 2D12, Abcam), or rabbit anti-β-actin pAb (1:5000, ab8227, Abcam). Blots were incubated with anti-mouse or anti-rabbit secondary antibodies conjugated to IRDye 680RD or IRDye 800CW, respectively, for 1 h at 23 °C (1:20,000; LI-COR Biosciences). Blots were quantified on the Odyssey scanner (LI-COR Biosciences).

### 2.5. Fluorescence data acquisition and analysis

Fluorescence lifetime (FLT) measurements were conducted in a top-read FLT plate reader designed and built by Fluorescence Innovations, Inc. (St. Paul, MN) in 384-well plate formats. GFP donor fluorescence was excited with a 473 nm microchip laser from Concepts Research Corporation (Belgium, WI), and emission was acquired with 490 nm long-pass and 520/17 nm band-pass filters (Semrock, Rochester, NY). We previously validated the performance of this FLT plate reader with known fluorescence standards, as well as with a FRET-based high-throughput screening strategy that that targeted 2-color SERCA.^28,29^ FLT waveforms from donor- and donor/acceptor-labeled samples were analyzed as described in our previous publications and the supplemental methods.^28,29^

### 2.6. Ca-ATPase activity assay

An enzyme-coupled, NADH-linked ATPase assay was used to measure Ca^2+^-activated ATPase activity of SERCA in 96-well microplates. Each well contained HEK293-6E homogenates, 50 mM MOPS (pH 7.0), 100 mM KCl, 1 mM EGTA, 0.2 mM NADH, 1 mM phosphoenol pyruvate, 10 IU/mL of pyruvate kinase, 10 IU/mL of lactate dehydrogenase, 3.5 μg/mL of the Ca^2+^ ionophore A23187, and CaCl_2_ added to set [Ca^2+^]_i_ to the desired values.^11^ The assay was started by addition of Mg-ATP at a final concentration of 5 mM, and well absorbances were read in a SpectraMax 384 Plus microplate spectrophotometer (Molecular Devices, Sunnyvale, CA). The Ca-ATPase assays were conducted over a range of [Ca^2+^]_i_, and the ATPase activities were fitted using the Hill function

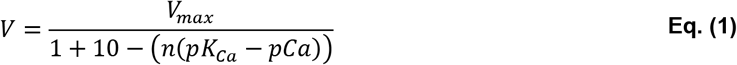

where V is the initial rate of Ca^2+^-dependent ATPase activity at a specified Ca^2+^ concentration (pCa), V_max_ is the rate of Ca-ATPase activity at saturating [Ca^2+^]_i_, n is the Hill coefficient, pK_Ca_ is the fitted Ca^2+^ dissociation constant, and pCa is the concentration of ionized Ca^2+^ per specific well and V.

### 2.7. Live-cell Ca^2+^ transient measurements

hiPSC-CMs (day 35 of differentiation) were enzymatically dissociated using the Detach 2 kit (Promocell, Heidelberg, Germany) and plated on Matrigel-coated German glass coverslips. After 2 days, the plated hiPSC-CMs were loaded with a Ca^2+^-sensitive fluorescent dye (Fura-2 AM, cell permeant, ThermoFisher, Rockville, MD, USA), and the ratio of fluorescence intensities (excitation ratio of 340/380 nm) were recorded using the IonOptix system (Ionoptix, Milton, MA). The electrically-induced Ca^2+^ transients were triggered by pulses from a MyoPacer (IonOptix, Milton, MA) at 40 V and 0.5 Hz, with cells at 23 °C. Ca^2+^ traces were analyzed using IonWizard software (IonOptix) to calculate the release amplitude (peak height relative to baseline) and tau (time of Ca^2+^ removal). The number of irregular Ca^2+^ transients was quantified using IonOptix software.

Using rAAV2.L31A, rAAV2.I40A, and rAAV2.L31A/I40A PLB viruses, hiPSC-CMs were infected at day 30 of differentiation (1e4 MOI), enzymatically dissociated on day 35, and plated on Matri-gel coated German glass coverslips, as described above. Fura-2 fluorescence measurements were recorded in AAV-infected hiPSC-CMs on day 37 and compared to non-infected control hiPSC-CMs. AAV viruses were purified according to the two-plasmid method with iodixanol gradient as described.^30^

### 2.8. Statistical analyses

All statistics were performed using Prism (GraphPad, La Jolla, CA), and analysis was done by one-way ANOVA followed by the Bonferroni post hoc test; analysis of two group comparisons was done by Student’s t-test (*P < 0.05 and ‘ns’ is not significant). Data are presented as mean ± standard error of the mean (SEM), and all statistical values were calculated from a minimum of three separate experiments.

## 3. Results

### 3.1. FRET assay demonstrates SERCA-binding competition between PLB_WT_ and loss-of-function PLB_M_

Previously, two classes of point mutations within the PLB TM domain had been identified via alanine-scanning mutagenesis, with disparate outcomes relative to SERCA: loss-of-inhibitory function (LOI) and gain-of-binding function (GOB).^8,31^ We hypothesized that combining LOI and GOB mutations in a single PLB would result in a LOI-PLB that can effectively compete with the inhibitory PLB_WT_. To test this, we used a SERCA2a-PLB biosensor system that consists of (a) co-expressed GFP-tagged SERCA2a and RFP-tagged PLB_WT_ in HEK293-6E cells and (b) a fluorescence lifetime (FLT) readout of FRET. FLT-FRET is used to resolve changes in SERCA2a-PLB complex structure and binding (Fig. 1A).^28^ We varied the ratio between the donor molecule (GFP-SERCA2a) and acceptor molecule (RFP-PLB_WT_), and found that the maximal energy transfer efficiency E (fractional decrease of the fluorescence lifetime) in this live-cell based system saturates at 0.10, as previously reported.^28^ We transfected cells expressing GFP-SERCA2a and RFP-PLB_WT_ with either untagged PLB_WT_ or PLB_M_ containing TM mutations (L31A, I40A, or L31A/I40A). Displacement of the RFP-PLBWT from its interaction with GFP-SERCA2a was observed as a decrease in the FRET efficiency relative to that measured in control cells expressing only the GFP-SERCA/RFP-PLB_WT_ donor-acceptor pair (Fig. 1B). FRET between GFP-SERCA2a and RFP-PLB_WT_ decreased significantly upon co-expression of unlabeled PLB_WT_, indicating that the RFP-PLB_WT_ and unlabeled PLB_WT_ are in equilibrium for hetero-dimeric binding to GFP-SERCA2a. An alanine substitution at L31A (LOI mutation) did not significantly alter the FRET value relative to the wild-type control. In contrast, an alanine substitution at I40A (associated with increased SERCA inhibition via decreased Ca^2+^ sensitivity) further decreased FRET, indicating that I40A-PLB_M_ competes effectively with RFP-PLB_WT_ with a potency (affinity) comparable to or greater than that of unlabeled PLB_WT_. The combination of L31A and I40A mutations resulted in a PLB_M_ with binding similar to that of I40A-PLB_M_, consistent with our hypothesis that PLB_M_ with both mutation types can retain high affinity toward SERCA2a.

Expression of the constructs was confirmed by SDS-PAGE and immunoblot (Fig. 1C, D). GFP-tagged SERCA2a was resolved from endogenous SERCA2b, as verified by binding of a GFP-specific antibody. For GFP-SERCA, there were no bands of lower mobility (apparent proteolysis) and no bands of higher mobility (apparent aggregation) in intact cells. Expression of RFP-PLB produced a single band recognized by RFP- and PLB-specific antibodies in samples heated to 95°C prior to electrophoresis. The I40A mutation disrupts pentamer formation, and we observed the disappearance of the band corresponding to PLB pentamer when the I40A mutation is present (Fig. 1D; lanes 11 and 12). PLB pentamer was detected for PLB_WT_ and L31A-PLB (Fig. 1D; lanes 13 and14). As the expression level of GFP-SERCA2a, RFP-PLB_WT_, and PLB_M_ is not significantly different between respective samples, we conclude that the observed changes in FRET are due to displacement of RFP-tagged PLB_WT_ via competition (to GFP-SERCA2a) with untagged PLB_M_. The Ca-ATPase activity of GFP-SERCA is similar to that of untagged SERCA, so the GFP tag does not perturb endogenous function.^25^

### 3.2. Effects of loss-of-function and gain-of-binding PLB mutants on SERCA2a regulation

The effects of PLB_M_ overexpression on the Ca^2+^-dependence of SERCA2a activity (measured by pK_Ca_) were determined in HEK293-6E homogenate samples, enabling co-expression of PLB_WT_ and PLB_M_ in a cell-based system similar to the FRET assays. Ca-ATPase activity in HEK293-6E cells expressing SERCA2a alone (Fig. 2A, black) increases with Ca^2+^ at physiological Ca^2+^ concentrations (*e.g.*, between pCa 7 and pCa 6) and saturates at higher Ca^2+^ concentrations (pCa 5), consistent with previous measurements.^28^ Concomitant expression of PLB_WT_ (Fig. 2A,B, blue) decreases Ca^2+^ affinity (increases pK_Ca_) of SERCA2a, and the additional expression of PLB_WT_ (purple) or L31A-PLB_M_ (cyan dashes) does not significantly decrease the apparent Ca^2+^ affinity; indicating that SERCA2a is fully inhibited and that L31A-PLB_M_ is not sufficient to relieve PLB_WT_ inhibition under these experimental conditions. We observed a further decrease in the apparent Ca^2+^ affinity upon co-expression of I40A-PLB_M_ (red) with SERCA2a and PLB_WT_, suggesting that I40A-PLB_M_ shows competitive binding to SERCA2a in the presence of PLB_WT_ (consistent with FRET measurements) and acts as a super-shifter/inhibitor.^8,11,32^ Co-expression with L31A/I40A-PLB_M_ (green) increased Ca^2+^ affinity to values similar to the SERCA-only sample, indicating that the PLB inhibition was relieved. Taken together, the structural (FRET) and functional (Ca-ATPase) assay results demonstrate that L31A/I40A-PLB_M_ is non-inhibitory and binds to SERCA2a in HEK293-6E cells.

**Fig. 2.**
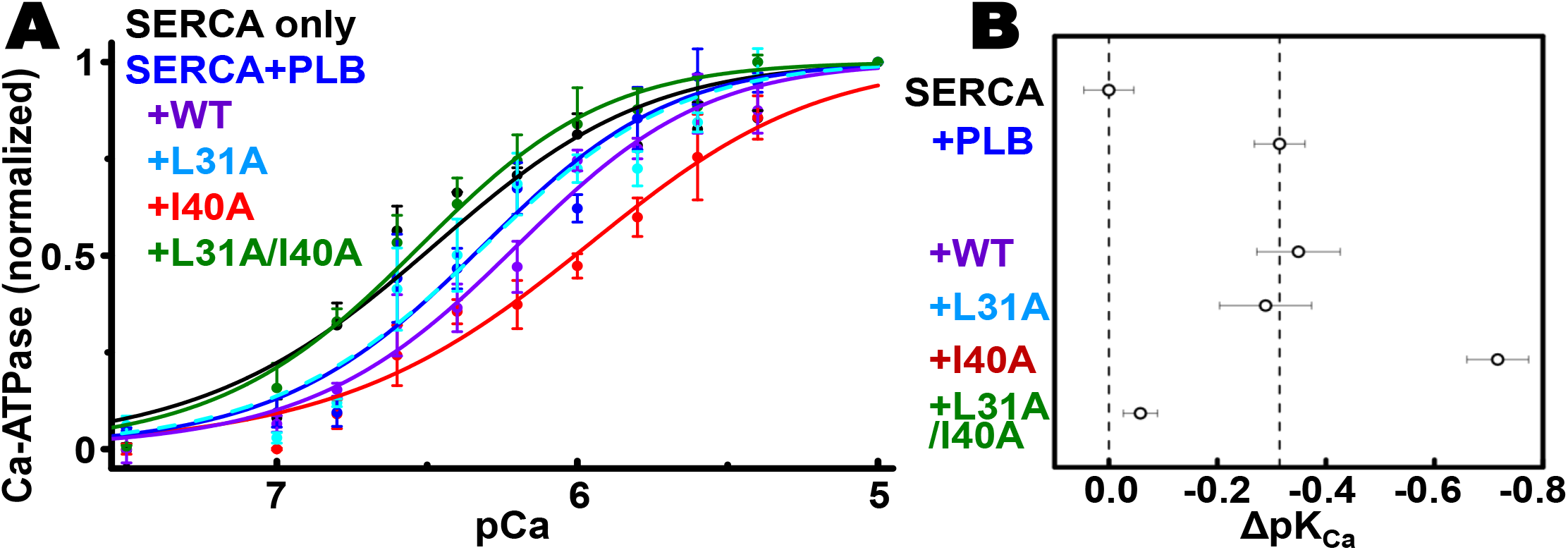
SERCA activation by PLB_M_ competition. (A) Normalized Ca-ATPase activity of HEK293-6E cell homogenates was measured 48 hours after transfection using +empty vector (negative control, not shown), +WT-SERCA2a (black), +WT-SERCA2a/WT-PLB (blue), +WT-SERCA2a/WT-PLB +WT-PLB (purple), +WT-SERCA2a/WT-PLB +L31A-PLB (cyan, dashed line), WT-SERCA2a/WT-PLB +I40A-PLB (red), or WT-SERCA2a/WT-PLB +L31A/I40A-PLB (green). (B) Quantification of apparent Ca^2+^ affinity (1/pK_Ca_) and PLB_M-_ induced change (Δ) from (A). Error bars indicate SEM (n = 3). ***Note to journal: this is a 2-column wide figure***.

### 3.3. rAAV-driven expression of L31A/I40A-PLB_M_ enhances Ca^2+^ transients in hiPSC-CMs

We differentiated hiPSCs derived from a healthy individual (SKiPS-31.3 line) into cardiomyocytes using established protocols that yields a predominant ventricular-like population.^27^ To assess physiological relevance in this native SERCA-PLB cardiac system, we tested the effects of rAAV-delivered PLB_M_ by infecting hiPSC-CMs with empty virus, rAAV2.L31A-, rAAV2.I40A-, or rAAV2.L31A/I40A-PLB_M_ at day 30 of differentiation and recording Ca^2+^ transients via fura-2AM upon electrical stimulation at day 37 (Fig. 3A). Ca^2+^ transients were regular in appearance (Fig. 3B). hiPSC-CMs expressing rAAV2.L31A-PLB_M_ or rAAV2.I40A-PLB_M_ did not show significant changes in peak amplitude or the rate of Ca^2+^ removal relative to control cells expressing empty vector (Fig. 3C). However, there were striking improvements in Ca^2+^ transient parameters (increased peak amplitude, increased Ca^2+^ removal rate) in hiPSC-CMs infected with rAAV2.L31A/I40A-PLB_M_ (Fig. 3B and C). Expression of the exogenous PLB_M_ was confirmed by qRT-PCR where virally-infected hiPSC-CMs had approximately two PLB_M_ per one PLB_WT_ (Fig. 3D). We observed no differences in SERCA2a levels indicating that enhanced Ca^2+^ transport is due to altered PLB regulation and not a compensatory change in SERCA2a expression (Fig. 3D). Cardiac troponin T expression levels were also unchanged, suggesting that PLB_M_ doesnot significantly alter hiPSC-CM differentiation.

**Fig. 3.**
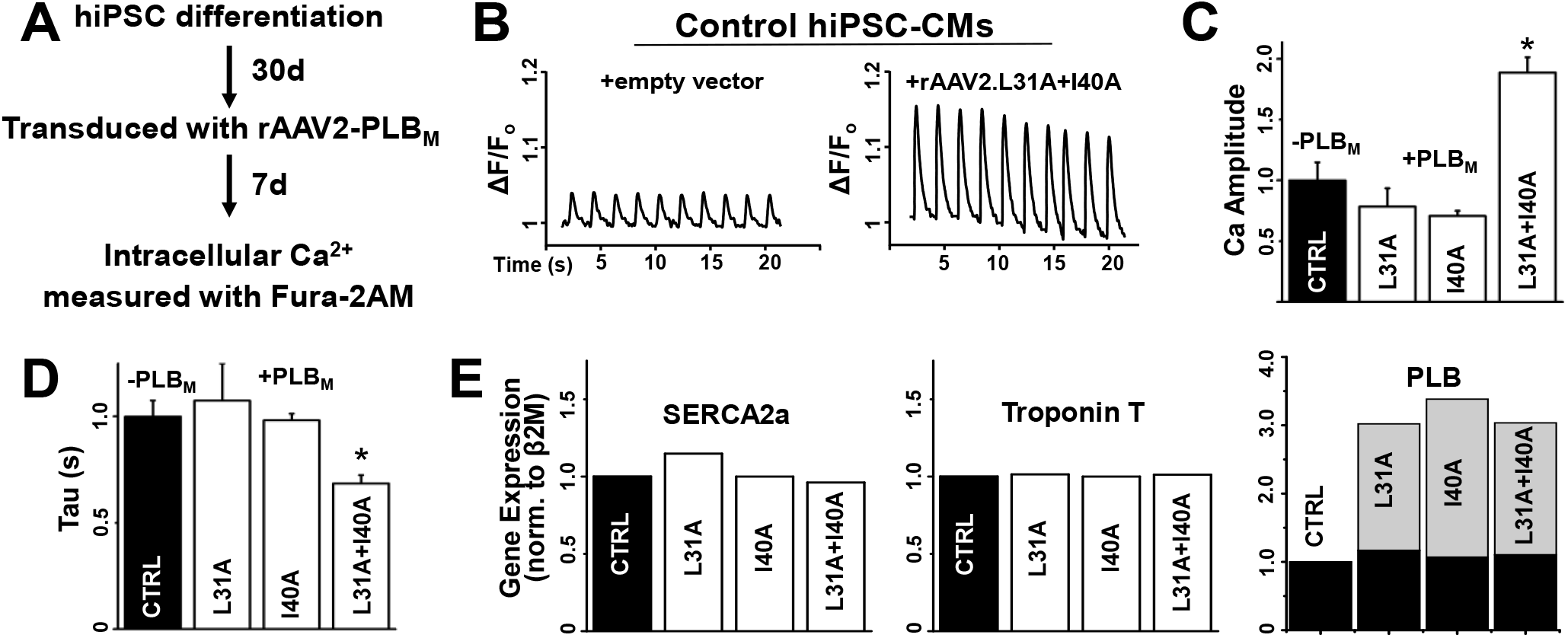
In normal, healthy hiPSC-CMs, rAAV2-driven expression of L31A/I40A-PLB enhances Ca^2+^ release amplitude and Ca^2+^ removal rate. (A) Schematic of experimental design for hiPSC differentiation, transduction, and cytosolic Ca^2+^ assay. (B) Representative Ca^2+^ transients in control hiPSC-CMs *(left)* compared to rAAV2-L31A/I40A PLB transduced hiPSC-CMs *(right)*. (C) Quantification of Ca^2+^ release amplitude (peak height) normalized to uninfected control (CTRL) and (D) Ca^2+^ removal time constant (Tau) of hiPSC-CMs normalized to uninfected control (CTRL). Error bars indicate SEM (n = 3). *P < 0.05. (E) qRT-PCR analysis of gene expression of Ca^2+^ transport and contractility proteins: SERCA2a *(left)*, cardiac troponin T *(center)*, and PLB *(right;* black = endogenous PLB_WT_, gray = exogenous PLB_M_) of uninfected (CTRL) or infected (rAAV2-L31A/I40A PLB) hiPSC-CMs. Gene expression values are normalized to uninfected control. ***Note to journal: this is a 2-column wide figure***.

### 3.4. Relief of SERCA2a inhibition by PLB_M_ rescues irregular Ca^2+^ transients in cardiomyopathic R14del-PLB hiPSC-CMs

Autosomal dominant mutations in PLB have been linked to DCM and these mutations include R9C^33^, R14del^34–36^, R25C^37^, and L39stop^38^. Recent work using hiPSC-CMs derived from patients heterozygous for the R14del-PLB mutation show mislocalization of R14del-PLB into aggregates leading to higher levels of autophagy and accompanying irregular Ca^2+^ transients.^39,40^ Mice heterozygous for the R14del-PLB mutation have severe defects in SERCA2a activity and Ca^2+^ transport. We sought to characterize the effects of viral expression of PLB_M_ in this dilated cardiomyopathic model to evaluate therapeutic potential. To accomplish this, we generated a R14del-PLB knock-in hiPSC line using homologous recombination via CRISPR/Cas9 to insert the R14del-PLB mutation into the control SKiPS-31.3 line (Fig. 3), producing an isogenic control to test our therapies.^27^ While the R14del-PLB hiPSC-CMs display no differences in Ca^2+^ transients parameters measured at day 21 (compared to wild type cells), we observed an arrhythmic (irregular) Ca^2+^ transient profile in R14del-PLB hiPSC-CMs after day 37 of differentiation. This irregularity occurred at a frequency of 58 ± 19% of Ca^2+^ transient recorded under paced conditions. (n = 85 cells) (Fig. 4A/D). This phenotype switch was also observed in patient-derived R14del hiPSC-CMs.^39^ Upon infection with rAAV2.L31A/I40A-PLB, we observed an improvement in Ca^2+^ handling properties of the R14del-PLB hiPSC-CMs. Representative Ca^2+^ tracings show a significant reduction in irregular Ca^2+^ transients and improvements in Ca^2+^ amplitude and tau upon rAAV2.L31A/I40A-PLB infection (Fig. 4B/C). Finally, no irregular Ca^2+^ transients (n = 20 cells) were observed after infection with rAAV2.L31A/I40A-PLB (Fig. 4D). Altogether, these data indicate an impairment in Ca^2+^ cycling in R14del-PLB hiPSC-CMs that is corrected by exogenous expression of the SERCA activating L31A/I40A-PLB_M_.

**Fig. 4.**
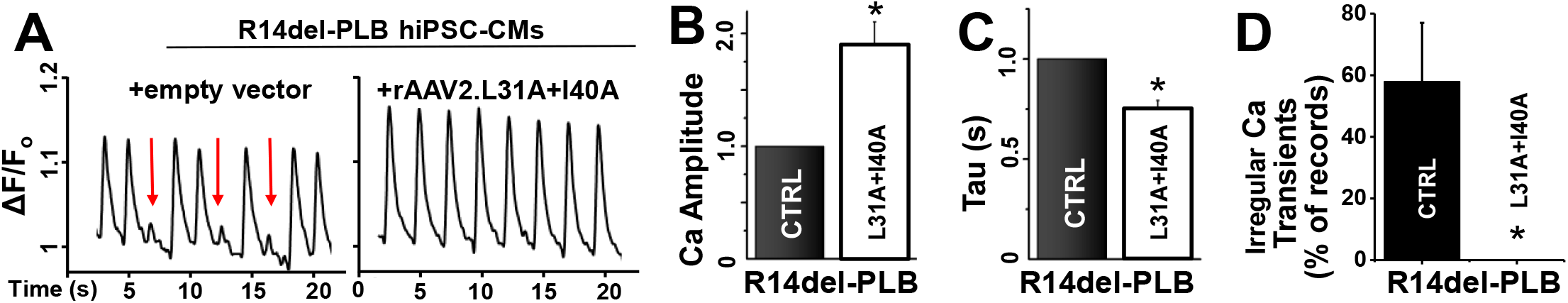
In cardiomyopathic R14del hiPSC-CMs, viral delivery of L31A/I40A-PLB rescues arrhythmogenic Ca^2+^ transients and enhances cellular Ca^2+^ cycling. (A) Representative Ca^2+^ transients in R14del PLB hiPSC-CMs (left) andR14del hiPSC-CMs infected with rAAV2.L31A/I40A-PLB (right). Arrhythmogenic Ca^2+^ transients are identified by red arrow. (B) Ca^2+^ release peak amplitude (fluorescence intensity) normalized to uninfected R14del hiPSC-CMs. (C) Ca^2+^ re-uptake time constant (Tau) normalized to uninfected R14del hiPSC-CMs. (D) Irregular Ca^2+^ transients determined by quantifying the number of individual cells exhibiting irregular Ca^2+^ transients. Error bars indicate SEM (n = 3). *P < 0.05. **Note to journal: This is a 2-column wide figure**.

## 4. Discussion

Hallmarks of heart failure include decreased contractile velocity, decreased relaxation rates, and pathological remodeling (*i.e.*, ventricular hypertrophy or dilation).^1^ Although the critical events that lead to impaired cardiac performance are still being determined, it is clear that pathways controlling intracellular Ca^2+^ homeostasis significantly contribute to decreased cardiomyocyte and contractile function.^2–4^ SERCA2a and PLB are major determinants of SR Ca^2+^ transport in the heart, and alterations to their function has profound effects on intracellular Ca^2+^ cycling.^5^ We demonstrated previously that the interaction between the cardiac Ca^2+^ pump, SERCA2a, and its principal inhibitor, PLB, can be measured via FRET in HEK293 cells.^28^ In the present study, we applied this FRET assay to measure the competition of non-inhibitory PLB_M_ and RFP-labeled PLB_WT_ for binding to GFP-SERCA2a. We used this assay to measure the relative affinity of non-inhibitory PLB_M_ and identified a double mutant (L31A/I40A) that binds with affinity greater than PLB_WT_. Expression of L31A/I40A-PLB_M_ increased Ca-ATPase activity in PLB_WT_-expressing HEK293 cells. Mechanistically, these results strongly suggest that L31A/I40A-PLB_M_ enhances SERCA activity by competitively displacing the inhibitory form of PLB. These results demonstrate that it is possible to separate the inhibitory potency of PLB from its binding affinity. Specifically, the properties of the PLB double mutant (L31A = loss-of-inhibition = LOI/I 40A = gain-of-binding = GOB) show the dominant effect of the LOI mutation (over the GOB mutation) while preserving binding affinity for SERCA2a.

There were previous attempts to increase SR Ca-influx in HF-animal models by expressing non-inhibitory PLB_M_, including a pseudo-phosphorylated mutant (S16E)^41^ and a dominant-negative mutant (R3E/R14E)^42^. Although there were significant improvements in Ca transients and cardiac output, these mutations preclude PLB phosphorylation by PKA and β-adrenergic stimulation.^41,42^ Expression of an unregulated PLB_M_ (*e.g.*, S16E) could cause chronic inotropic stimulation, whereas the PLB_M_ double mutant L31A/I40A remains phosphorylatable at the S16 site. In hiPSC-CMs, we found that rAAV2-driven expression of L31A/I40A-PLB_M_ significantly improved Ca^2+^ release amplitude and Ca^2+^ removal in both healthy and pathogenic cell lines. Using CRISPR/Cas9 we created a knock-in hiPSC cell line that is heterozygous for the DCM-causing mutation R14del-PLB. We differentiated this cell line into cardiomyocytes to study potential defects in Ca transport and homeostasis. Similar to hiPSC-CMs derived from R14del-PLB human patients,^39^ our cell line developed an arrhythmia-like Ca^2+^ transient phenotype by day 37 of differentiation. We demonstrated that expression of L31A/I40A-PLB_M_ reversed Ca transport dysfunction and irregular Ca^2+^ transients in R14del-PLB hiPSC-CMs. These results may justify future studies to test potential therapeutic effects *in vivo*. Altogether, this work paves the way for a potential therapeutic for heart diseases associated with impaired Ca^2+^ transport and decreased contractility.

## Abbreviations

Ca^2+^: (calcium)
B2M: (β2-microglobulin)
DCM: (dilated cardiomyopathy)
ER: (endoplasmic reticulum)
FLT: (fluorescence lifetime)
FRET: (fluorescence resonance energy transfer)
GOB: (gain of binding function)
GFP: (green fluorescent protein)
hiPSC-CM: (human induced pluripotent stem cell-derived cardiomyocyte)
HEK: (human embryonic kidney)
HF: (heart failure)
LOI: (loss of inhibitory function)
mAb: (monoclonal antibody)
miRNA: (microRNA)
pAb: (polyclonal antibody)
PLB: (phospholamban)
PLB_M_: (phospholamban mutant)
PLB_WT_: (wild type phospholamban)
rAAV: (recombinant adeno-associated virus)
RFP: (red fluorescent protein)
SERCA2a: (sarcoendoplasmic reticulum calcium ATPase 2a)
SR: (sarcoplasmic reticulum)

## Acknowledgments

This study was supported by N.I.H. grants to D.D.T. (GM27906, HL129814, and AG26160). The content is solely the responsibility of the authors and does not necessarily represent the official views of the National Institutes of Health. We thank Bengt Svensson for help with figures. Spectrophotometric assays were performed in the Biophysical Technology Center at the University of Minnesota Department of Biochemistry, Molecular Biology, and Biophysics.

## Supplementary Methods

### 1.1. Fluorescence Data Acquisition and Analysis^28^

The measured FLT waveform

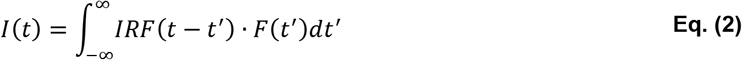

is a function of the nanosecond decay time t, and is modeled as the convolution integral of the measured instrument response function, IRF(t), and the fluorescence decay model, *F*(t). The fluorescence decay model

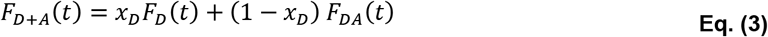

is a linear combination of a donor-only fluorescence decay function *F*_D_(t) and an energy transfer-decreased donor fluorescence decay *F*_DA_(t). The donor decay *F*_D_(t) is a sum of *n* exponentials

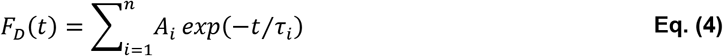

with discrete FLT species τ_i_ and pre-exponential mole fractions *A*_i_. For the GFP donor, two exponentials (*n* = 2) are required to fit the observed fluorescence decay waveform. The energy transfer-decreased donor decay function, *F*_DA_(t),

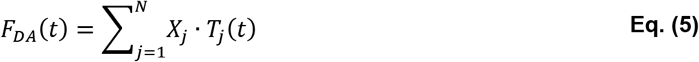

is the sum of the distribution of multiple structural states (*j*) with mole fractions *X*_j_, represented by the FRET-quenched donor fluorescence decays *T*_j_(t). The increase in the donor decay rate (inverse donor FLT) due to FRET is given by the Förster equation

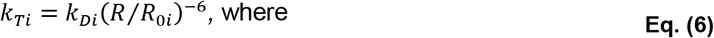

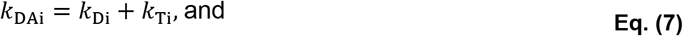

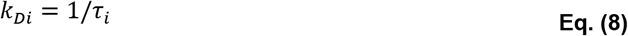

**Supplementary Table 1.**
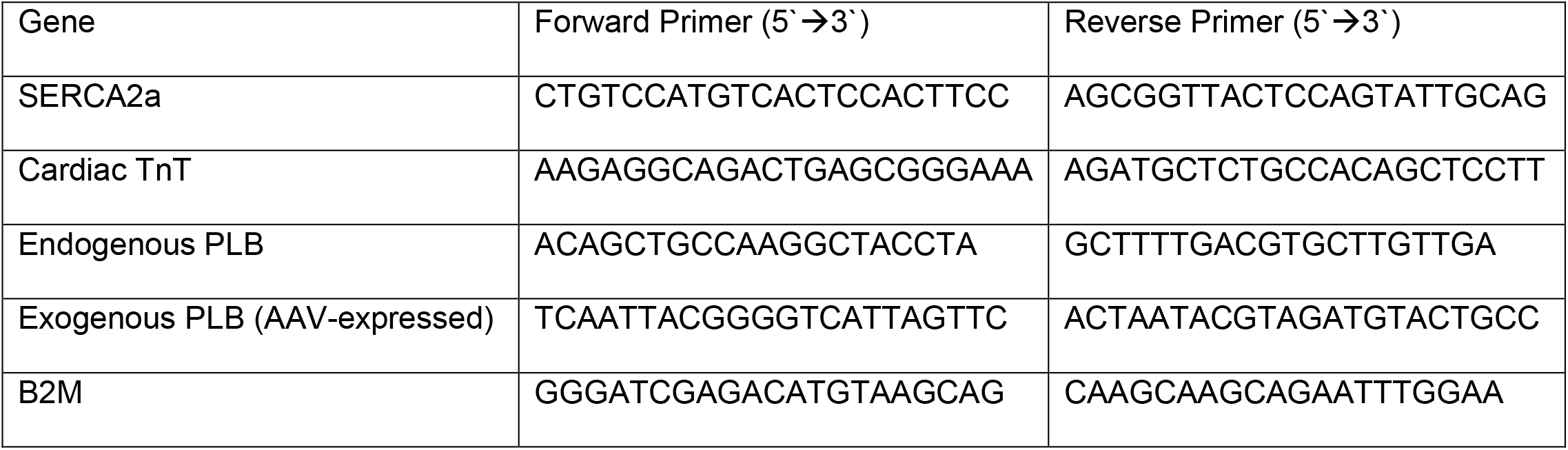
Primers used in qRT-PCR.

